# General auditory and speech-specific contributions to cortical envelope tracking revealed using auditory chimeras

**DOI:** 10.1101/2020.10.21.348557

**Authors:** Kevin D. Prinsloo, Edmund C. Lalor

## Abstract

1.

In recent years research on natural speech processing has benefited from recognizing that low frequency cortical activity tracks the amplitude envelope of natural speech. However, it remains unclear to what extent this tracking reflects speech-specific processing beyond the analysis of the stimulus acoustics. In the present study, we aimed to disentangle contributions to cortical envelope tracking that reflect general acoustic processing from those that are functionally related to processing speech. To do so, we recorded EEG from subjects as they listened to “auditory chimeras” – stimuli comprised of the temporal fine structure (TFS) of one speech stimulus modulated by the amplitude envelope (ENV) of another speech stimulus. By varying the number of frequency bands used in making the chimeras, we obtained some control over which speech stimulus was recognized by the listener. No matter which stimulus was recognized, envelope tracking was always strongest for the ENV stimulus, indicating a dominant contribution from acoustic processing. However, there was also a positive relationship between intelligibility and the tracking of the perceived speech, indicating a contribution from speech specific processing. These findings were supported by a follow-up analysis that assessed envelope tracking as a function of the (estimated) output of the cochlea rather than the original stimuli used in creating the chimeras. Finally, we sought to isolate the speech-specific contribution to envelope tracking using forward encoding models and found that indices of phonetic feature processing tracked reliably with intelligibility. Together these results show that cortical speech tracking is dominated by acoustic processing, but also reflects speech-specific processing.

This work was supported by a Career Development Award from Science Foundation Ireland (CDA/15/3316) and a grant from the National Institute on Deafness and Other Communication Disorders (DC016297). The authors thank Dr. Aaron Nidiffer, Dr. Aisling O’Sullivan, Thomas Stoll and Lauren Szymula for assistance with data collection, and Dr. Nathaniel Zuk, Dr. Aaron Nidiffer, Dr. Aisling O’Sullivan for helpful comments on this manuscript.

**Significance Statement:** Activity in auditory cortex is known to dynamically track the energy fluctuations, or amplitude envelope, of speech. Measures of this tracking are now widely used in research on hearing and language and have had a substantial influence on theories of how auditory cortex parses and processes speech. But, how much of this speech tracking is actually driven by speech-specific processing rather than general acoustic processing is unclear, limiting its interpretability and its usefulness. Here, by merging two speech stimuli together to form so-called auditory chimeras, we show that EEG tracking of the speech envelope is dominated by acoustic processing, but also reflects linguistic analysis. This has important implications for theories of cortical speech tracking and for using measures of that tracking in applied research.

## 3. Introduction

Over the past few years research on natural speech processing has benefited from recognizing that low frequency cortical activity tracks the amplitude envelope of natural speech (Ahissar et al., 2001; Lalor & Foxe, 2010; Huan Luo & Poeppel, 2007). This has been useful for investigating the mechanisms underlying speech processing (Peelle & Davis, 2012), how such processing is affected by attention (Nai Ding & Simon, 2012; Power, Foxe, Forde, Reilly, & Lalor, 2012; E. M. Zion-Golumbic et al., 2013), and how audio and visual speech interact (M. J. Crosse, Butler, & Lalor, 2015; M. J. Crosse, Di Liberto, & Lalor, 2016; H. Luo, Liu, & Poeppel, 2010; Zion-Golumbic, Cogan, Schroeder, & Poeppel, 2013). However, it remains unclear to what extent these cortical measures reflect higher-level speech-specific processing versus lower-level processing of the spectrotemporal/acoustic stimulus dynamics.

There has been some evidence that speech intelligibility affects these envelope tracking measures (Jonathan E Peelle, Joachim Gross, & Matthew H Davis, 2013), suggesting that they may indeed index speech-specific processing. But precisely what aspects of speech processing are reflected in envelope tracking measures, or even how specifically the measures reflect speech processing at all, is unclear. It has been suggested that different neural populations, having different functional roles in receptive speech processing, may simultaneously contribute to envelope tracking measures (Nai Ding & Simon, 2014). Furthermore, specific mechanistic theories have been proposed, suggesting that envelope tracking (or envelope “entrainment” more specifically) represents a more active process for parsing speech into discrete chunks for further processing (Giraud & Poeppel, 2012). However, drawing definitive inferences about the meaning of cortical speech tracking must contend with the likelihood that much of the speech-tracking signal will derive from general auditory processing of the stimulus acoustics by cortical regions that are agnostic to the special nature of speech. Indeed a wealth of evidence has amassed suggesting that speech is processed by a hierarchically organized network of cortical regions with responses in earlier stages (including primary auditory cortex) being well accounted for based on the spectrotemporal acoustics of the stimulus, and later stages being invariant to those acoustics and involved in more abstract linguistic processing (Davis & Johnsrude, 2003; de Heer, Huth, Griffiths, Gallant, & Theunissen, 2017; DeWitt & Rauschecker, 2012; Huth, de Heer, Griffiths, Theunissen, & Gallant, 2016; Kell, Yamins, Shook, Norman-Haignere, & McDermott, 2018; Norman-Haignere & McDermott, 2018). This is consistent with the idea that speech sounds are perceived using mechanisms that evolved to process environmental sounds more generally (Diehl, Lotto, & Holt, 2004), with additional linguistic processing occurring in specialized downstream pathways (Hickok & Poeppel, 2007; Rauschecker & Scott, 2009).

Indeed, this notion that cortical tracking of speech might reflect (perhaps a lot of) general acoustic processing as well as (perhaps a more limited contribution from) linguistic processing helps to explain several other findings in the literature. For example, while cortical envelope tracking sometimes shows sensitivity to speech intelligibility as mentioned above (Jonathan E Peelle et al., 2013), this is definitely not always the case (Howard & Poeppel, 2010). Indeed robust cortical tracking has been observed for completely unintelligible speech, including vocoded and backward speech (Di Liberto, Crosse, & Lalor, 2018; Di Liberto, O’Sullivan, & Lalor, 2015; Howard & Poeppel, 2010), as well as very general auditory stimuli such as amplitude modulated broadband noise (Lalor, Power, Reilly, & Foxe, 2009). So acoustic processing definitely makes a substantial contribution. In the present study, we aim to explore this idea of dissociable contributions to envelope tracking using, so-called, auditory chimeras (Smith, Delgutte, & Oxenham, 2002). In particular, we record EEG from subjects as they listen to speech-speech chimeras – stimuli comprised of the temporal fine structure (TFS) of one speech stimulus modulated by the amplitude envelope (ENV) of another speech stimulus. By varying how these chimeras are constructed, we obtain some control over which stimulus is recognized by the listener, allowing us to decouple the processing of the acoustic envelope from the speech content. We hypothesize that envelope tracking will be dominated by the dynamic changes in the acoustic energy of the signal, with a smaller component reflecting speech-specific processing.

## 4. Materials and Methods

### Subjects

Seventeen native English speakers (mean age 24.9; std 3.7; range 20-30; 8 males) participated in the experiment. Participants reported no neurological diseases, and self-reported normal hearing. Informed consent was obtained from all participants before the experiment and subjects received monetary compensation for their time. The study was conducted in accordance with protocols approved by the Research Subjects Review Board at the University of Rochester.

### Stimuli and Experimental Procedure

In our experiment, we wanted to decouple the acoustic amplitude fluctuations of the stimuli from their speech content. To do this, we used so-called auditory chimeras (Smith et al., 2002). These are stimuli in which the envelope of one sound is used to modulate the temporal fine structure of a second sound. Importantly, this can be done after first filtering the sounds into complementary frequency bands using a filter bank. Then, each filter output is Hilbert transformed to derive its analytic signal, and the envelope (calculated as the magnitude of the analytic signal) of the first sound is used to modulate the fine structure (calculated as the cosine of the phase of the analytic signal) of the other sound within each band – giving a series of partial chimeras. Finally, the partial chimeras are summed over all frequency bands to produce the final chimera (Fig. 1). Critically for our experiment, the number of frequency bands used has a marked effect on which original sound source is actually perceived (Smith et al., 2002). If the original sounds are both speech and just one band is used, then listeners will partially recognize the speech content corresponding to the temporal fine structure. They will (obviously) fail to understand any of the speech content corresponding to the source of the broadband envelope. But, as the number of frequency bands grows, listeners will increasingly understand the speech content relating to the source of the envelope and will no longer perceive the speech content of the temporal fine structure source.

**Figure 1.**
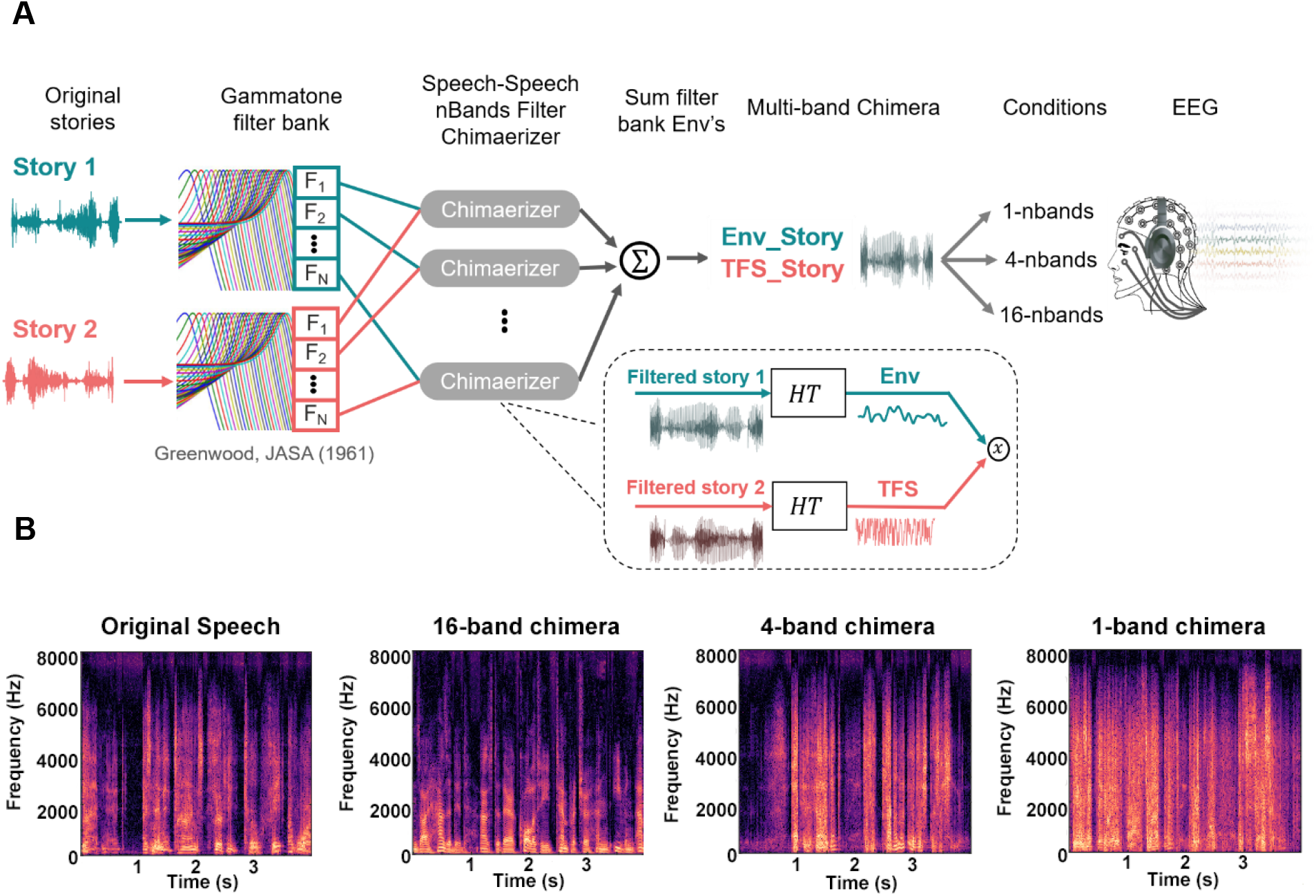
Speech-speech chimera generation. **A**. Two audiobooks were passed through Gammatone filterbanks with different numbers of frequency bands (1, 4, 16) for our different experimental conditions. The outputs of these filterbanks were Hilbert transformed (HT) allowing us to calculate the envelope of story 1 and a temporal fine structure representation of story 2. The envelope of story 1 was used to modulate the temporal fine structure of story 2 within each frequency band, and the resulting partial chimeras were summed to produce a final multi-band chimera that was played to the subject. **B**. Spectrogram of an example segment of story 1 (left), and of the three types of chimera corresponding to that segment.

For our experiment, we generated auditory chimeras from two speech sources. These were audiobooks of two classic works of fiction (Env_Story and TFS_Story) which were read in English by two different male speakers and sampled at 48 kHz. Our study consisted of three experimental conditions each involving the presentation of a speech-speech chimera generated using a filterbank with a different number of frequency bands, specifically 1, 4, and 16. These filterbanks were comprised of FIR band-pass filters that were spectrally divided into logarithmically spaced filters along the cochlea frequency map from 80 to 8020 Hz (Greenwood, 1990). Each filter had a nearly rectangular response, and adjacent filters overlapped by 25% of the bandwidth of the narrower filter. All stimulus processing was performed using MATLAB 2016.b software (The MathWorks Inc.). And stimuli were presented to subjects using the Psychophysics toolbox (Brainard & Vision, 1997) within the MATLAB environment along with custom code.

Forty-five separate one-minute speech segments were randomly selected from each audiobook and were used to generate the three types of chimera (1, 4, and 16 bands), resulting in 15 trials per chimeric condition. All of the chimera stimuli were made by using the envelope of the Env_Story segment to modulate the TFS of the TFS_Story segments. We chose to always use one audiobook as the Env_Story and the other audiobook as the TFS_Story. This was mostly driven by a desired not to divide our EEG data into an extra set of subconditions. We also wanted to be consistent in the behavioral task in having subjects always answering on one story for the envelope, and always the other story for TFS. Importantly, we used these same two audiobooks in previous studies (Power et al., 2012; O’Sullivan et al. 2015), and saw no systematic difference in the strength of neural tracking to each story. As such, we think it is unlikely that we will have introduced any bias into our results by choosing to not counterbalance the audiobooks across the Env and TFS conditions. All stimuli were spatialized by convolution with a head-related transfer function (HRTF), simulating a scenario in which each stimulus appears to be spatially located directly in front of the subject (Algazi, Duda, Thompson, & Avendano, 2001). Incidentally, we selected an additional ten segments from Env_Story and five segments from TFS_Story, and we presented these – unmodified – to the subjects as a control condition. Ultimately, however, we did not include any analysis of these control data in our results below.

Subjects were instructed to attend to the audio stimulus and maintain visual fixation for the duration of each trial on a crosshair centered on the screen, and to minimize eye blinking and all other motor activities. To quantify speech intelligibility, after each trial, subjects were required to answer four multiple-choice questions (MCQs) on both stories (i.e., 4 from the Env_Story and 4 from the TFS_Story). Each question had four possible answers. MCQs, answer choices and chimera condition order were all pseudorandomized across subjects. Stimulus presentation and data recording took place in an audiometric grade sound attenuated and electromagnetically shielded room (IAC Acoustics, North Aurora, IL. 120A-Series). The visual stimuli (crosshair, MCQs and answer choices) were presented on a 24-inch LCD monitor (ASUS Predator), operating at a refresh rate of 60 Hz, and participants were seated at a distance of 70 cm from the display. All audio stimuli were normalized to have the same root mean square intensity and were presented binaurally through Sennheiser HD650 headphones at a self-adjusted comfortable level.

### EEG Acquisition and Preprocessing

EEG was recorded from 130 channels at 512 Hz using a BioSemi ActiveTwo system. 128 cephalic electrodes were positioned according to the BioSemi Equiradial system, with another 2 electrodes located over the left and right mastoids. Triggers indicating the start of each trial were presented using Psychophysics toolbox in MATLAB for synchronous recording along with the EEG.

The EEG data were first resampled to 128 Hz using the *decimate* function in MATLAB. The *decimate* function incorporates an 8^th^ order low-pass Chebyshev Type I infinite impulse response (IIR) anti-aliasing filter. Consistent with previous research suggesting that speech tracking is strongest in the δ-band (1-4 Hz) and θ-band (4-8 Hz), we focused our analysis on frequencies below 8 Hz. Specifically, we used a zero phase-shift Chebyshev type-2 bandpass filter with pass bands between 1 and 8 Hz. Subsequent preprocessing was performed using the Fieldtrip toolbox (Oostenveld, Fries, Maris, & Schoffelen, 2011) and custom code in MATLAB. After filtering, bad channels were defined as those whose variance was either less than half or greater than twice that of the surrounding 3–7 channels (depending on location in the montage). These channels were then replaced through spherical spline interpolation (Fieldtrip). Next after removing bad channels we applied denoising using independent component analysis, usually only removing one or two components reflecting eye-movement-related artifacts, which were determined following definitions provided in (Debener, 2010).

### Indexing Cortical Speech Tracking using th e Temporal Response Function

The goal of this study was to examine how cortical activity tracks the envelope of speech and how that tracking might derive from acoustic vs speech-specific processing. To index cortical speech tracking, we used the temporal response function framework (Michael J Crosse, Di Liberto, Bednar, & Lalor, 2016). The general idea of this framework is to use linear regression to map between ongoing speech features (e.g., the envelope) and ongoing neural responses. This can be done either in the forward direction, by examining how well different speech features can explain variance in EEG responses on individual channels (a forward encoding model approach). Or it can be done in the backward direction by attempting to reconstruct an estimate of a speech feature using all of the EEG channels (a decoding approach). It provides a number of dependent measures: the accuracy of EEG predictions based on a forward encoding model, the accuracy of stimulus reconstructions based on a backward decoding model, or the weights that are applied to the stimulus features (forward) or EEG data (backward) on different channels and at different timelags between stimulus and response (see (Michael J Crosse et al., 2016) for more details). In the forward direction, the TRF can be described via the following equation:

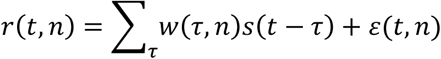

where *r*(*t, n*) is the neural response at timepoint *t* on channel *n, s*(*t*) is the stimulus feature at time *t*, which can be a univariate (e.g., the envelope) or multivariate (e.g., the spectrogram) representation of the speech, *τ* indexes the relative time lag between the speech stimulus feature and the neural response in samples, and *ε*(*t, n*) is an error term. In our analysis, *t* runs from 1 to the length of the trial (i.e., 60 s), *n* = 1 … 128 channels, and *τ* = –100 to 500 ms indicating that we are exploring the impact of the stimulus on the EEG data at lags from –100 ms to +500 ms. We then estimated the unknown TRF, *w*(*τ, n*) using regularized (ridge) linear regression (Michael J Crosse et al., 2016). As mentioned above, this enabled us to use the weights of the TRF itself (*w*(*τ, n*)) as dependent measures, and to test how well different speech features were represented in the EEG by seeing how well the TRF can predict EEG responses to held out trials (Michael J Crosse et al., 2016). We also conducted a backward decoding analysis described by the following equation:

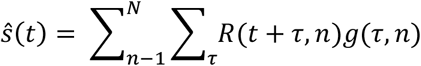

where the main difference is that the TRF, *g*(*τ, n*), is a multivariate function (often known as a decoder) that was fit on all EEG channels at the same time. In general, this approach is more sensitive as it makes more effective use of the available data, but it is also limited in terms of what speech features, *Ŝ* (*t*), can be reconstructed. In our case, we restricted ourselves to decoding based on the univariate amplitude envelope of the speech stimuli (please see next section). Again, the weights of the decoder and the decoder’s ability to reconstruct the stimulus were then available to us as dependent measures.

### Speech stimulus representations

To use the TRF framework to examine how cortical activity tracks the amplitude envelope of our speech chimeras, we first needed to calculate the amplitude envelope of our stimuli. Moreover, we were specifically interested in how this cortical tracking might reflect contributions from acoustic and speech-specific processing. To do this, we also wished to use the TRF to explore how cortical activity reflects the processing of other acoustic and speech-specific features of our stimuli. Importantly, for other acoustic and speech-specific features to show up in measures of envelope tracking, their temporal dynamics would need to correlate with those of the envelope. This is true for the spectrogram and phonemes of speech (and less true for higher level representations of speech based on semantic content, e.g., Broderick et al., 2018). As such, to assess how contributions from spectrotemporal and phoneme processing might contribute to envelope tracking, we derived the following acoustic and phonemic feature representations of our speech stimuli:

1. *The Envelope (Env)*. We calculated the amplitude envelope for each of the 45 one-minute speech segments from both Env_Story and TFS_Story. We did this by first bandpass filtering the segments into 128 logarithmically-spaced frequency bands spanning 80 and 8000 Hz using a cochlear filterbank (Greenwood, 1990). The envelope for each band was computed using the Hilbert transform, then the broadband envelope was obtained by averaging over the 128 narrowband envelopes. The output of this process was then logarithmically transformed in an effort to account for the nonlinear relationship between electrophysiological responses and stimulus amplitude (Aiken and Picton, 2008). Specifically, dB envelope representations were generated by taking 20log_10_ of the broadband envelope (Aiken and Picton, 2008).
2. *Recovered-envelope (RE-Env)*. One important issue with modeling our EEG data as a function of the envelopes of the original stories used in creating the chimeras, is that the subjects were not actually presented with these original stories. So, when considering how our EEG might track the envelope of the stories in a chimera, we wanted to understand how the cortex might be able to recover the envelope of the TFS story (TFS_Story) from the output of the cochlea. To estimate this recovered envelope, we did the following. We first determined the TFS of the chimera stimuli for all three chimera conditions (1, 4, and 16 bands) by calculating the Hilbert transform of those chimera stimuli, and then determining the cosine of the phase of the resulting analytic signal. We then filtered the TFS signal into 128 bands using a cochlear frequency map spanning a range of 80 to 8020 Hz (Greenwood, 1990) and we created analytical signals for each frequency band. Finally, the envelope of each of these narrow bands (calculated as the absolute values of its analytical signal) were summed to generate the broadband *RE-Env* (Smith et al., 2002; Zeng et al., 2004).
3. *Spectrogram (Sgram)*. The envelope is a very impoverished measure of a speech signal. To more fully explore the relationship between envelope tracking and the acoustic processing of speech we sought to more richly represent the speech acoustics. We did this by computing the Log-Mel spectrogram of our speech stimuli (Chi, Ru, Shamma, & KWG, 2005; Schädler, Meyer, Kollmeier, & CYE, 2012; Verhulst, Altoè, & Vasilkov, 2018). This involved passing the speech signals through a bank of 64 filters that spanned from 124.1 to 7284.1 Hz and that were organized according to the Mel scale (i.e., the scale of pitches judged by listeners to be equal in distance one from another). The output of these filters was then scaled by a logarithmic compressive nonlinearity to convert it to the final Log-Mel spectrogram. The choice of Log-Mel spectrogram was made because it incorporates several properties of the auditory system, non-linear frequency scaling and compression of amplitude values.
4. *Phonemes (Ph)*. We were also interested to explore how envelope tracking might relate to speech-specific processing in the form of a sensitivity to phonemes within the speech. To derive a representation of phonemes, we used the Montreal Forced Aligner (McAuliffe, Socolof, Mihuc, & Wagner…, 2017), a python-based open source tool based the Kaldi ASR toolkit that relies on triphone-based Hidden Markov Models to create statistical models associating phonetic symbols to speech signals (http://kaldi-asr.org/). The aligner, given an audio speech file and corresponding textual orthographical transcription, partitions each word into phonemes from the American English International Phonetic Alphabet (IPA) and performs forced-alignment (Yuan and Liberman, 2008), returning the starting and ending time-points for each phoneme. This information was then converted into a multivariate time-series composed of indicator variables, which are binary arrays (one for each phoneme). These are active for the time-points in which phonemes occurred. The phonemes are mutually exclusive, so that only one can be active at each sample point. We selected a subset of the IPA comprised of the 35 most frequent phonemes in the presented speech stimuli (3 of 38 IPA phonemes were excluded as being outliers in terms of how rare they were). *Ph* is a language dependent representation of speech.

### Using the m TRF to Assess the Sensitivity of EEG to Different Speech Features

In the results section that follows, we will be using the TRF framework to assess how well different speech features are represented in the EEG data. As discussed above, two of our dependent measures will be: 1) how well we can reconstruct a speech envelope from the EEG responses (i.e., backward modeling); and, 2) how well we can predict data on different EEG channels using the stimulus spectrogram and phonemes (i.e., forward modeling). Two key related considerations for using these dependent measures are: 1) how to train and test the mTRF models so as to trust in the generalizability of the findings, and 2) how to regularize the mTRF models so as to make them robust to EEG fluctuations that are unrelated to the speech, as well as to noise. In terms of the latter issue, we used ridge regression. In brief, this approach penalizes large values in the mTRF, meaning that we are reducing the variance of our mTRF weights, by adding a bias term. Ultimately, this makes for a more generalizable model (see Crosse et al., 2016 for extensive discussion). However, care must be taken not to over-regularize. In what follows we describe how we determined the ridge regression parameters (known as lambda values) that allowed us to best map between speech features and EEG.

Our general strategy for training, regularizing, and testing was to use the following cross-validation procedure. For each of our four stimulus representations, a separate TRF was fit to each of M trials for a range of lambda values. One trial was ‘left out’ to be used as a ‘test set’, with the remaining M-1 trials to be used for the inner cross-validation. Next one of these inner M-1 trials was chosen to be ‘left out’ and used as a ‘validation set’. The remaining M-2 trials were used as a ‘training set’. An average model was obtained by averaging over the single-trial models in the ‘training set’. This was done for each lambda value. Next, this average model was used to either reconstruct the speech envelope by convolving it with the EEG data (decoding) or predict the EEG responses by convolving it with the chosen speech representations (forward modeling) associated with the ‘validation set’. The accuracy of this reconstruction or prediction (of selected EEG channels, please see below) was assessed by comparing it with the real speech envelope or EEG using Pearson’s correlation coefficient. This procedure was repeated such that each of the M-1 trials was ‘left out’ of the ‘training set’ once. The lambda value that produced the highest average reconstruction or prediction accuracy across all of the validation sets was then chosen as the optimal lambda. Please note, this could mean different optimal values of lambda for reconstruction, for prediction, for each subject, and for each set of speech features.

Next, using the optimal lambda value chosen above, another average model was obtained by averaging over the single-trial models in both the ‘validation’ and ‘training’ sets and using that model on the data from the ‘test set’. Model performance was assessed by quantifying how accurately the reconstructed envelope or predicted EEG correlated with the actual stimulus envelope or the actual recorded EEG response from the ‘test set’, again using Pearson’s r. This entire procedure was repeated M times such that each trial was ‘left out’ of the inner-cross validation procedure once. As before the overall model performance was then finally assessed by averaging the overall individual model performances for each trial. Again, lambda parameter optimization was done separately for each stimulus representation and subject (i.e., each model was based on its respective optimal performance). Our modeling procedures were subjected to permutation testing to quantify a null distribution. 95% quartiles, demarcated by grey boxes, are reported in the figures.

To evaluate whether either phonetic or spectral acoustic features contributed independently to predicting the neural responses across conditions, we computed the partial correlation coefficients (Pearson’s r) between the EEG predicted by either phonological (or spectral-acoustic) feature model with the actual recorded EEG after controlling for the effects of spectral-acoustics (or phonological features). Specifically, we fitted separate cross-validated forward model TRFs on each of the two speech representations (Sgram, Ph) and predicted EEG based on those models. Then we used the built-in MATLAB function partialcorr (X, Y, Z) where X = actual recorded EEG, Y = predicted EEG in response to the feature of interest (the feature whose unique contribution is to be identified) and Z = concatenated predicted EEGs in response to the other feature (feature that is to be partialled out). This function computes the partial correlation coefficients between X and Y, while controlling for the variables in Z (Fisher 1924).

### Statistics

Significance at the group level of either decoder or encoder accuracies were evaluated using nonparametric permutation statistics. The neural responses were permuted across trials such that they were matched to features from a different trial, and the same leave-one-out cross-validation procedure as described above was performed to compute TRFs and either reconstruction or prediction accuracies. This was done 2,000 times for each subject to establish a distribution of chance-level prediction accuracies. By randomizing the data across trials and recalculating the test statistic, we obtained a reference distribution to evaluate the statistic of the actual data. We also ran randomization using cluster-based statistics (Monte Carlo procedure) across channels and time for topographical demarcation of significant time-sensor clusters to be used as regions of interest (ROI) for further analyses (Maris & Oostenveld, 2007; R. Oostenveld & Maris, 2007). Furthermore, we adopted the same procedures for the partial correlation analyses, based on the prediction accuracies (phonetic/spectral acoustic encoders), we also computed a distribution of partial correlations for the phonetic (or spectral acoustic) features measures (i.e. we partialled out the contributions of all other features). Unless otherwise stated, further analyses were done on significant channels following permutation tests or computed on 12 temporal parietal channels (6 symmetric pairs over left and right hemisphere).

Linear mixed-effects models (LME) were implemented to explore behavioral and EEG results and their interrelationship, via the *fitlme* function in MATLAB using the restricted maximum likelihood (REML) method. Advantages over standard analysis of variance (ANOVA) approaches have been previously reported (Krueger & Tian, 2004; Luke, 2017; Wainwright, Leatherdale, & Dubin, 2007). After visual inspection of the residual plots, it was clear there were no obvious deviation from homoscedasticity or normality. All *p*-values were estimated using the Satterthwaite approximations. Post-hoc analyses were performed using linear hypothesis testing on linear regression model coefficients (*coeftest*). Mixed-effects models account for multiple comparisons. Subjects were treated as random factors according to the following linear-model expression: (*LME* = (*RA*_*data*_ ∼ 1 + *Beh*_*data*_ + (1|*Subjects_ID*)) and (*LME* = (*RA*_*data*_ ∼ 1 + *Cond*_*nBands*_ + (1|*Subjects_ID*)), where *RA* stands for reconstruction accuracy and *Beh* corresponds to behavioral task performance.

As well as using frequentist probability-based statistics, we also used the Bayesian analog of an ANOVA (*anovanBF*) to allow us to explicitly determine the amount of evidence in favor of the null hypothesis (*H*_0:_ no interaction). We estimated the Bayes factors (*BF*_10_) using Matlab code adapted from RStudio (R-Core-Team, 2016; the function *anovanBF* in the toolbox *Bayes factor* (Morey, Rouder, & Jamil, 2015)). We adopted the commonly used Jeffrey-Zellner-Siow (JZS) prior with a scaling factor of 0.707 (Rouder, Morey, Speckman, & Province, 2012; Rouder, Speckman, Sun, Morey, & Iverson, 2009; Schönbrodt, Wagenmakers, Zehetleitner, & Perugini, 2017). Monte-Carlo resampling with 10^6^ iterations was used for the *BF*_10_ estimation. Subjects represented the random factor. Importantly, this estimation allows us to quantify evidence that our experimental factors and interactions explain variance in the data above the random between-subject variations. Standard convention stipulates that any *BF*_10_ exceeding 3 is evidence in favour of the alternative hypothesis (*H*_1_), while below 0.33 is in support of the null hypothesis (*H*_0_), and *BF*_10_ ranging between 1-3 is taken as weak anecdotal evidence in favor of the alternative hypothesis, as is widely reported in the literature (Wagenmakers, Wetzels, Borsboom, & Van Der Maas, 2011).

## 5. Results

128-channel EEG was recorded from seventeen subjects as they listened to 60-s auditory chimeras composed of segments of narrative speech from two audiobooks. In particular, the envelope of Env_Story was used to modulate the temporal fine structure of TFS_Story. By varying the number of frequency bands used in making the chimeras (1, 4, or 16 bands), we aimed to manipulate which story was understood by the subjects. Following Smith et al. (2002), we expected subjects to (partially) recognize TFS_Story for the 1-band chimeras, to (partially) recognize Env_Story for the 4-band chimeras, and to clearly recognize Env_Story for the 16-band chimeras.

With such a pattern of behavioral results, what might we expect in terms of variation in how cortical signals track the envelopes of the Env_Story and the TFS_Story? Based on previous studies, several plausible alternative hypotheses can be proposed as to how such signals might vary across our chimera conditions. We briefly suggest present four alternative hypotheses that could be plausibly advanced based on previous research.

Hypothesis 1 (Fig. 2*A*) is that neural tracking of speech is overwhelmingly dominated by acoustic processing. Were this true, one might expect the acoustic energy fluctuations (i.e., the envelope) to play such a dominant role that neural tracking would reflect the processing of only the Env_Story across chimera conditions. An example of a previous study that might motivate a hypothesis like this is Howard & Poeppel (2010), which reported that discrimination of speech stimuli based on neuronal response phase patterns depends on acoustics but not comprehension. Hypothesis 2 (Fig. 2*B*) is that neural tracking of speech is dominated by speech-specific processing. Were this hypothesis true, one might expect neural tracking to very closely mirror speech intelligibility, with no tracking for the unintelligible stimulus despite its acoustic energy fluctuations (e.g., no tracking of the Env_Story in the 1-band condition). This would be a reasonable hypothesis based on a series of studies by Zoefel and VanRullen who explored speech entrainment using novel speech/noise speech stimuli that were constructed to have no systematic changes in sound amplitude and spectral content (Zoefel & VanRullen, 2016; Zoefel & VanRullen, 2015) It might also be a reasonable hypothesis coming from studies that have focused on how speech-specific acoustic “edges” might be driving neural entrainment to that speech (Doelling, Arnal, Ghitza, & Poeppel, 2014). Hypothesis 3 (Fig. 2*C*) is that neural tracking of speech is dominated by general acoustic processing that can be enhanced by intelligibility through either predictive processing related to the speech content (e.g., Broderick et al., 2019), by attention (O’Sullivan et al., 2015), or higher order statistical structure in the acoustics (e.g., Zuk et al., 2020). Were this hypothesis true, one might expect that neural tracking would again be dominated by the acoustic energy of the Env_Story, but that the tracking of Env_Story would increase across chimera conditions. One would expect no discernible tracking of the TFS_Story in this case. Finally, hypothesis 4 (Fig. 2*D*) is that neural tracking of speech contains contributions from low-level neural populations that are responsive to general acoustic input and from higher-level neural populations that are specifically tuned to speech sounds. Were this hypothesis true, one would expect the Env_Story to contribute to all chimera conditions, but more so in chimera conditions were the Env_Story is intelligible. And one would expect the TFS_Story to contribute only in those conditions where the TFS_Story is intelligible. This is the hypothesis we favour based on previous literature (and a synthesis thereof). If the data support this hypothesis, it will also be of interest to see the relative size of the contributions to the EEG that appear to derive from general acoustic processing vs speech-specific processing.

**Figure 2.**
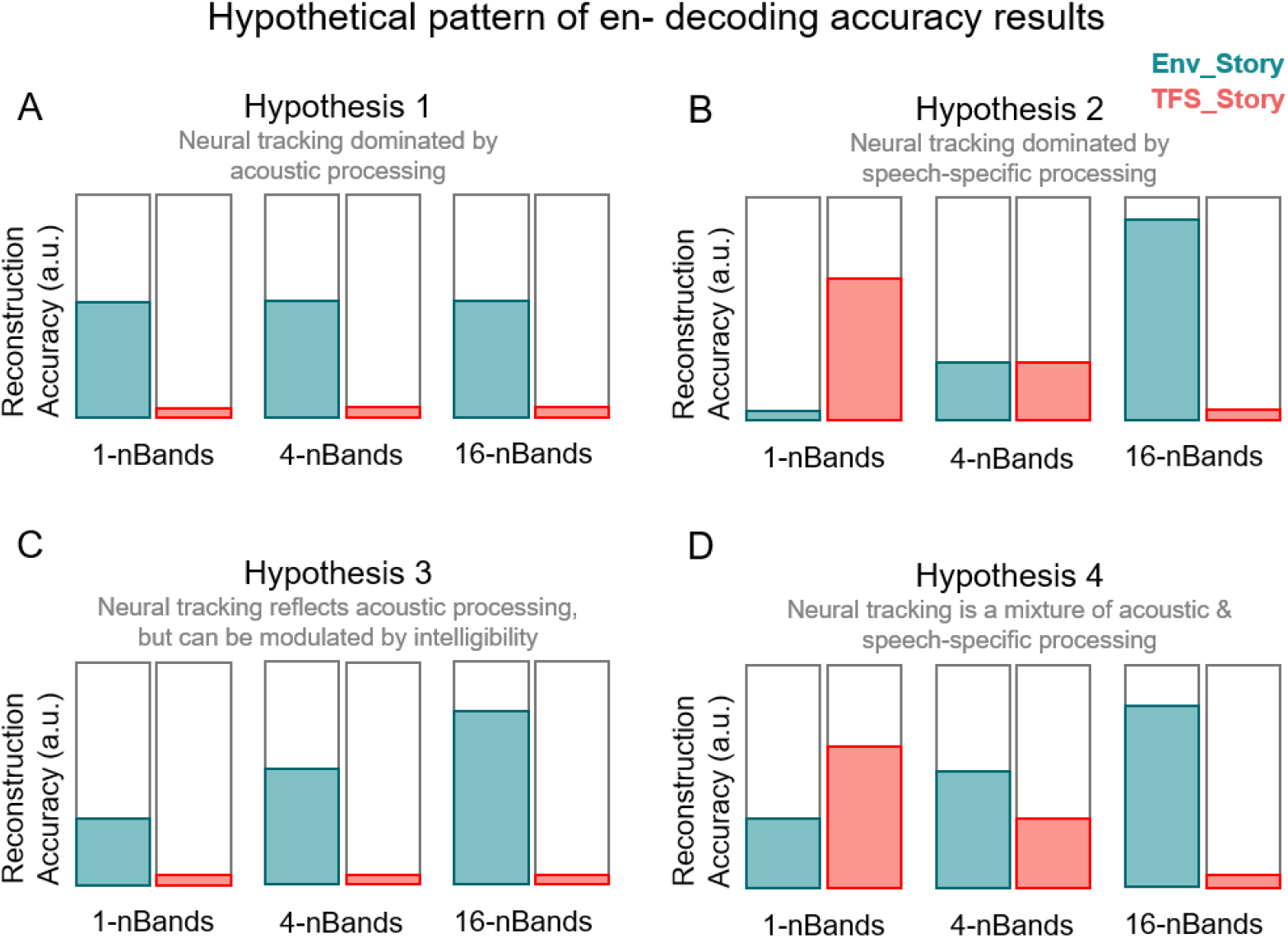
Hypothetical pattern of results. **A**. Decoding accuracy is mainly driven by the speech envelope and is not modulated as a function of speech intelligibility. Temporal fine structure (TFS) is poorly decoded as there is no envelope feature. **B**. Speech is the dominant factor here, neural tracking is speech specific and is related to the parsing of the structure of higher order speech features. In this instance, it would be predicted that there is stronger tracking for the most intelligible conditions, i.e. 1-band for the TFS_Story and 16-bands for the Env_Story. The lack of tracking of an unintelligible stimulus component that varies in its acoustic energy could be explained by comodulation masking release (CMR), wherein coherent fluctuations in a masker can improve signal detection by suppressing that masker (Dau, Ewert, & Oxenham, 2009). **C**. The hypothesis that obligatory sensory tracking of the acoustics of speech stimulus is enhanced by speech intelligibility. Tracking is dominated by the speech envelope as shown in the Env_Story, while there is little to no cortical tracking of the TFS_Story. **D**. Neural tracking is a mixture of both general acoustic processing and speech-specific processing. In this case, we would expect to see tracking of the Env_Story across all conditions, with stronger tracking as that story becomes more intelligible. We would also expect to see some tracking of the TFS_Story in conditions where that story is intelligible. How strongly the tracking of the Env_Story will be relative to the TFS_Story is an open question.

### Speech comprehension varies across speech-speech chimera conditions

Speech comprehension was assessed by asking multiple-choice questions at the end of each trial. There were four possible answers to each question setting the theoretical chance level at 25%. However, we determined that, at the group level, a score of 27.5% was significantly greater than chance (*p* = 0.05) based on a binomial distribution using all 840 trials (14 subjects x 60 questions) per condition. We then tested our distribution of subject scores against 27.5% using a one-sided Wilcoxon sign-rank test. The pattern of behavior was largely consistent with what we expected based on Smith et al., 2002 (Fig. 3*A*). In particular, performance on the questions for Env_Story in the 1-band condition was 29.43% ± 2.11 (% mean ± SEM) and was not significantly greater than chance (p = 0.28). However, as the number of chimera bands increased, so did performance with subjects achieving 33.43% ± 1.57 and 68.43% ± 2.09 for the 4-bands and 16-band conditions, respectively, both of which were significantly greater than chance (*z* = 3.30, *p* = 4.56 × 10^−04^, and *z* = 3.60, *p* = 1.58 ×10 ^−04^ Wilcoxon signed-rank test). Meanwhile, the performance on TFS_Story showed largely the opposite effect. Performance on the questions were 37.86% ± 1.41, 33.71% ± 1.16, and 22.79% ± 1.73 for the 1-band, 4-bands, and 16-bands conditions respectively. Importantly, these scores were significantly above chance for the 1-band (*z* = 3.24, *p* = 6.04 ×10 ^−04^) and 4-bands conditions (*z* = 3.11, *p* = 9.37 × 10^−04^). However, as expected, they were not significantly above chance for the 16-band condition (*z* = −2.52, *p* = 0.99). We wanted to more explicitly test for the interaction effect between chimera condition and story that we expected from Smith et al., (2002). To do this we used a two-way repeated measures ANOVA (*rm*ANOVA, with factors of *chimera condition: 1-, 4-, 16-band*, and *story: ENV Env_Story, TFS TFS_Story*). This analysis showed main effects of *chimera condition* (*F*_(1,2)_ = 42.39, *p* < 0.001, η^2^ = 0.15, *BF*_10_ = 3.259), and *story* (*F*_(1,1)_ = 189.42, *p* < 0.001, η^2^ = 0.21, *BF*_10_ = 2.131 ×10 ^18^). Importantly, it also revealed a significant interaction between *chimera condition* and *story* (*F*_(1,1)_ = 195.45, *p* < 0.001, η^2^ = 0.51, *BF*_10_ = 2.128 ×10 ^31^). Post hoc analyses were carried out using Bonferroni-corrected pairwise *t*-tests which showed that Env_Story intelligibility scores in the 16-band condition were significantly higher than those in the 1-band condition (*t*_(16)_ = 16.71, *p* < 0.001, *BF*_10_ = 1.4210 ×10 ^10^), and TFS_Story scores in the 1-band were significantly higher than 16-bands (*t*_(16)_ = 13.04, *p* < 0.001, *BF*_10_ = 1.5114 ×10 ^6^). This pattern of results generally replicates the findings from Smith et al. (2002), even though we assessed speech intelligibility by means of comprehension questions, while Smith et al. (2002) probed word recall.

**Figure 3.**
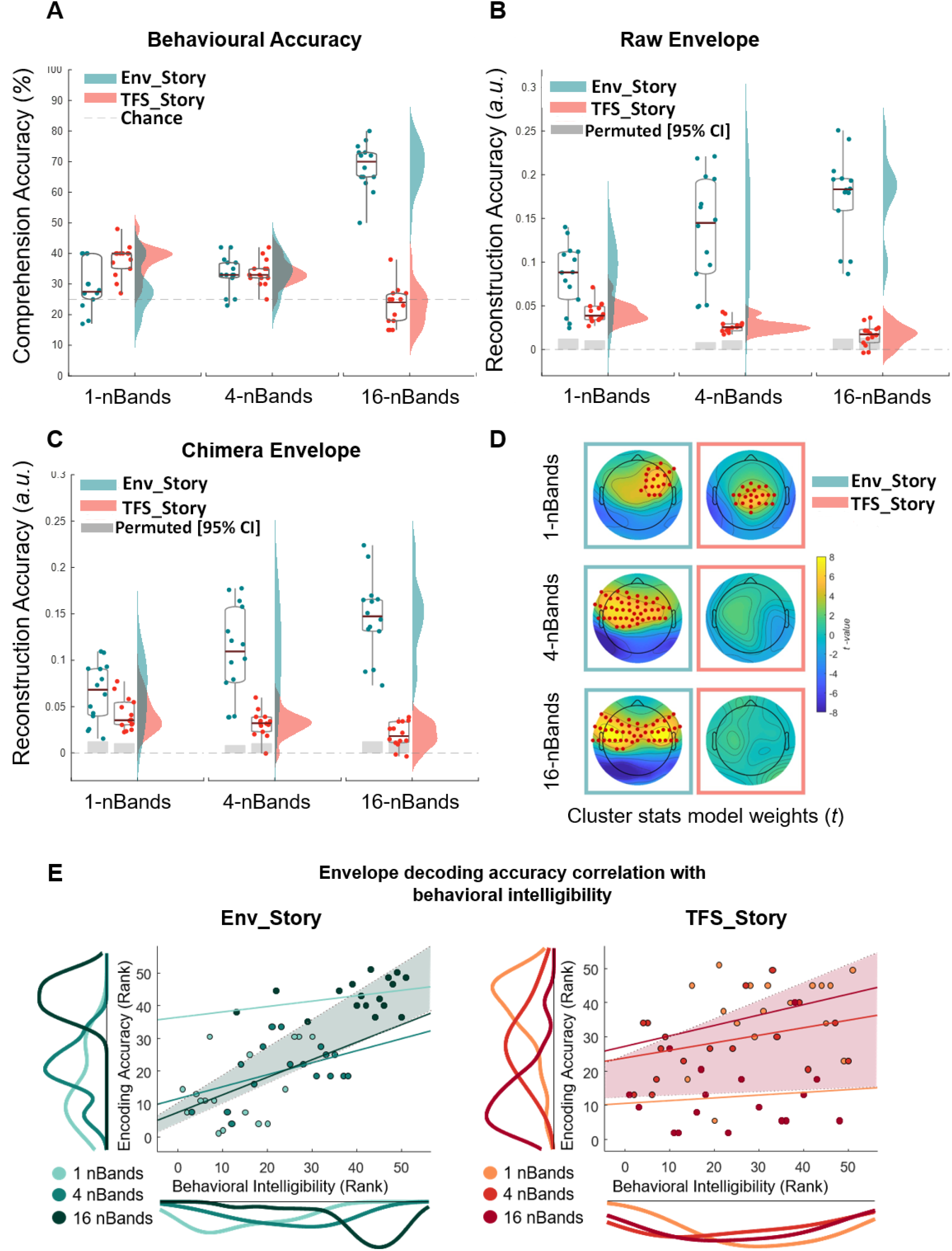
Cortical tracking of speech envelopes reflects both acoustics and speech intelligibility across chimera conditions. A-C Box plots (mean ± SEM) and rain kernel density estimates of speech intelligibility and reconstruction decoding accuracy (Rho) values for Env_Story (turquoise) and TFS_Story (peach). Group- and individual statistics; black line in box plot indicates mean across subjects, asterisks indicate group statistical significance (permutation test, *p < 0.05) and grey box demarcates single subject-level statistical significance above chance (permutation test, *p < 0.05). Single dots represent single subject data. **A**. Behavioral intelligibility, **B**. original envelope reconstruction accuracies, and **C**. chimera envelope reconstruction accuracies across bands, conditions, and stories. **D**. Topographical plots show forward transformed decoder weights across all channels for Env_Story (C) and TFS_Story (D) over a time-lag window of 80-120 ms. Red dots indicate significant effects at the group level (one-tailed cluster-based permutation test, N=2000, p < 0.05). **E. Left** Env_Story and **E. Right**. TFS_Story brain-behavior correlations. Individual dots represent subjects and are color coded according to condition. Correlations were assessed using robust Pearson’s correlation (bootstrap permutation test p < 0.05. This was done for each individual condition (colored lines), as well as collapsed across all conditions (shaded area representing confidence interval 95%). Subplots show the distribution of the data in terms of both envelope decoding (left) and behavior (below) using the same color code as the subject dots.

### Cortical tracking of the original story envelopes varies with understanding of speech-speech chimeras

To adjudicate between the four hypotheses introduced above (Fig. 2), and to test hypothesis 4 in particular, we wished to explore how cortical tracking of speech envelopes might reflect acoustic and speech-specific processing. Importantly, all three of our stimulus conditions (1-, 4-, and 16-band chimeras) have envelopes that are derived from Env_Story. As such, their acoustic energy will be dominated by that of Env_Story. However, as shown in the previous section, for the 1-band condition, subjects are more likely to understand the speech content from TFS_Story. How might envelope tracking reflect these acoustic and behavioral inconsistencies? We investigated this by attempting to reconstruct estimates of the speech envelopes corresponding to Env_Story and TFS_Story from the EEG recorded during each of our chimera conditions. As discussed above, we did this for all 15 trials for each condition using a cross-validation training and testing procedure. Importantly, for this first analysis, we simply sought to reconstruct the envelopes of the original speech segments, not of the chimeras themselves. Then we compared these reconstructions to the two original envelopes using Pearson’s correlation. We also determined a baseline level of reconstruction performance by generating a null distribution of Pearson’s r values by shuffling the labels between trials 1000 times and attempting to reconstruct the envelopes using the same procedure (grey bars demarcate 95% confidence interval).

The general pattern of stimulus reconstruction measures mirrored that of the behavioral results, and supported hypothesis 4 (Fig. 2*D*). Specifically, as the number of frequency bands in the chimeras increased, envelope reconstructions for Env_Story tended to increase and reconstructions for TFS_Story decreased (Fig. 3*B*). We assessed this pattern of results using a two-way rmANOVA (with factors of *chimera condition* and *story*). As with the behavioral results, we found significant main effects for *chimera condition* (*F*_(1,2)_ = 30.45, *p* < 0.001, η^2^ = 0.007, *BF*_10_ = 16.042) and *story* (*F*_(1,2)_ = 34.44, *p* < 0.001, η^2^ = 0.39, *BF*_10_ = 4.287). And, mirroring our key behavioral result, we found a significant interaction between *chimera condition* and *story* (*F*_(1,1)_ = 56.65, *p* < 0.001, η^2^ = 0.15, *BF*_10_ = 19.307). Again, we carried out post hoc analyses using Bonferroni-corrected pairwise *t*-tests. These also tracked with the behavioral results in that reconstructions for Env_Story were significantly higher for the 16-band condition than 1-band condition (*t*_(16)_ = 16.0, *p* < 0.001, *BF*_10_ = 6.01 ×10^8^), and TFS_Story reconstructions were significantly higher for the 1-band condition than the 16-band condition (*t*_(16)_ = 5.04, *p* < 0.001, *BF*_10_ = 270.01).

One important discrepancy between the behavioral and cortical tracking results is that there is robust cortical tracking of Env_Story for the 1-band condition (blue cloud, left, Fig. 3*B*), despite it being entirely unintelligible (blue cloud, left, Fig. 3*A*). Indeed, the cortical tracking of Env_Story in this condition is significantly stronger than that for TFS_Story (*t*_(16)_ = 5.25, *p* < 0.001, *BF*_10_ = 3.49 ×10^2^), despite TFS_Story being the one that is better understood. Of course, this is easily explained by reminding ourselves that the envelope of the chimera in this condition is determined by Env_Story. This highlights that fluctuations in the acoustic energy of an auditory stimulus will robustly drive cortical tracking, completely independently of any speech-specific activity. That said, the similar pattern of interaction effects for behavior and envelope tracking also supports the notion that cortical envelope tracking does, at least to some extent, index speech-specific processing. Env_Story tracking improved across conditions as Env_Story understanding increased, and TFS_Story tracking decreased as TFS_Story understanding fell. On this issue, it is particularly interesting to consider the case of the cortical tracking of TFS_Story in the 1-band condition. Again, the envelope of the chimera stimulus in this condition is dominated by Env_Story, and, yet, we have significant cortical tracking of the envelope of TFS_Story. How might this come about? We examine this further in the next subsection below where we will discuss the results of Fig 3*C*.

Before that, to visualize which EEG channels were contributing to the pattern of envelope reconstructions reported above, we transformed the backwards decoding model eights (Haufe et al., 2014) into a pattern of forward model weights (Fig. 3*D*). Visualizing these weights across the scalp is more analogous to exploring the topographical distribution of event-related potentials (Lalor et al., 2009). Using a cluster-based non-parametric permutation (*N* = 2000) analysis (Maris & Oostenveld, 2007) we identified model weights that were significantly different from zero across subjects and trials over a window of 80-120 ms time-lags. We chose these time-lags based on a prominent peak in the forward TRF model over these time-lags (please see below). For Env_Story there was an increase in the number of channels with significant model weights as intelligibility increased across conditions. These channels were largely over central and temporal scalp. For TFS_Story, significant model weights were found only for the 1-band condition, i.e., the condition for which TFS_Story was most intelligible. They were centered over central scalp regions.

We also sought to explore the correspondence in the pattern of results between behavior and stimulus reconstruction more directly (Fig. 3*E,F*). Specifically, we used robust correlation analysis with bootstrap resampling (Pernet, Wilcox, & Rousselet, 2013) to check for correlations between these measures across subjects. We did so in one analysis that included all conditions and found a strong positive correlation between decoding accuracy and behavioral intelligibility scores as a function of *chimera condition*, for Env_Story (*r*_*s*_ = 0.76, *p* < 0.001, 95% *CI* [0.61 0.85]) and TFS_Story (*r*_*s*_ = 0.31, *p* < 0.05, 95% *CI* [0.04 0.53]. We tested this relationship further using a linear-mixed effect model with subjects as a random factor (*LME* = (*RA*_*data*_ ∼ 1 + *Beh*_*data*_ + (1|*Subjects_ID*)) with comprehension accuracy scores and envelope reconstruction values as dependent and predictor variables respectively. The LME models variability due to stimulus conditions and subjects simultaneously. We found a significant positive relationship between reconstruction accuracy and comprehension scores for Env_Story (*β* = 415.73, *SE* = 43.91 *p* < 0.001) and for TFS_Story (*β* = 174.92, *SE* = 82.95 *p* < 0.05).

An important aspect of the previous result is that, as mentioned, it was carried out across all stimulus conditions collapsed together. We also wished to explore the possibility that there might be a correlation between behavior and reconstruction accuracy within each condition, across subjects. That said, we were not especially confident about finding such a result. This is because EEG envelope tracking measures can vary greatly between subjects based on factors that are completely unrelated to their ability to understand speech. For example, basic biophysical differences in cortical folding and brain/skull conductivity properties likely drive a very large percentage of the variance in EEG responses across subjects. With n = 17 in the present study, we expected such causes of variance might likely obscure any variance related to speech intelligibility across subjects. Indeed, using permutation testing while controlling for multiple comparisons across conditions, we found that there were no significant correlations within any one condition alone, for either the Env_Story (*p* = 0.28; *p* = 0.25; *p* = 0.21, for 1-bands, 4-bands, and 16 bands respectively) or the TFS_Story (*p* = 0.34; *p* = 0.27; *p* = 0.81, for 1-bands, 4-bands, and 16 bands respectively).

### Cortical tracking of envelopes recovered from the chimeras also varies with understanding of speech-speech chimeras

In the previous section, we wondered about how we might see significant cortical tracking of the TFS_Story envelope to a 1-band chimera whose envelope is determined by Env_Story (Fig. 3*B*). The answer likely lies in the fact each subject’s auditory cortex does not actually operate on the chimera stimuli that we present. Rather, each stimulus first undergoes extensive peripheral and subcortical processing. Indeed, the first stage of this processing is the fine-grained filtering by the cochlea. This produces a representation in the early auditory system that will be exquisitely sensitive to the temporal fine structure in the stimulus. It may be that some of the computations carried out by the cortex serve to synthesize a coherent TFS_Story speech object from these earlier representations – a process sometimes referred to as analysis-by-synthesis (Bever & Poeppel, 2010; N. Ding, Chatterjee, & Simon, 2013; Halle & Stevens, 1962; Halle, Stevens, Wathen-Dunn, & Woods, 1959; Poeppel & Monahan, 2011). This synthesis of an auditory object for TFS_Story from the temporal fine structure of the chimera is likely what our cortical envelope tracking measure is indexing, in this case.

To explore this more directly, we conducted an additional analysis aimed at examining how cortical activity might track with the envelopes of Env_Story and TFS_Story that we recovered from the actual chimera stimuli that were presented to the subjects. As detailed in the methods above, we determined the envelope of the chimera by passing it through a filter bank, determining the Hilbert transform of each narrowband frequency, and averaging the envelopes across frequency bands. Next, we computed the recovered envelope (ReEnv) of the TFS component from the chimeras using an approach based on the Hilbert transform and cochlear scaled filtering (Apoux, Millman, Viemeister, Brown, & Bacon, 2011; Irino & Patterson, 1997; Roy D Patterson, 1987; Roy D. Patterson, Uppenkamp, Johnsrude, & Griffiths, 2002). The overall pattern of results for the chimera envelopes (which should be very similar to those for Env_Story) and Re-Env (which we expected to resemble the envelopes for TFS_Story) was very similar to that for the original envelopes (Fig. 3*C*). Specifically, a two-way rmANOVA (with factors *chimera condition* and *story*) showed main effects for *chimera condition* (*F*_(1,2)_ = 29.05, *p* < 0.001, η^2^ = 0.006, *BF*_10_ = 15.002) and *story* (*F*_(1,1)_ = 32.41, *p* < 0.001, η^2^ = 0.32, *BF*_10_ = 4.024). And there was a significant two-way interaction between *chimera condition* and *story* (*F*_(2,32)_ = 55.45, *p* < 0.001, η^2^ = 0.14, *BF*_10_ = 19.007). Post hoc Bonferroni-corrected t-tests showed that Env_Story reconstruction accuracies for the 16-band condition were significantly higher than those for the 1-band condition (*t*_(16)_ = 16.84, *p* < 0.001, *BF*_10_ = 5.87 *x*10^8^), and TFS_Story reconstruction accuracies for the 1-band condition were significantly higher than that for the 16-band condition (*t*_(16)_ = 5.04, *p* < 0.001, *BF*_10_ = 280.21).

For Env_Story, the fact that the pattern of stimulus reconstruction results was so similar for the original and recovered envelopes (compare turquoise raincloud plots in Fig. 3*B* and 3*C*) was entirely unsurprising. This is because the envelopes from the original Env_Story were used to create the chimeras in the first place. As such, the raw stimulus envelope for the Env_Story and the envelope of the chimeras were highly correlated for all conditions (Table 1 – left column). For the same reason, the correlation between the raw stimulus envelope for the TFS_Story and the envelope of the chimeras was very low (in fact, it was slightly negatively correlated). Meanwhile, the similar pattern of stimulus reconstruction results for the original and recovered envelopes for TFS_Story (compare peach raincloud plots in Fig. 3*B* and 3*C*), was more interesting. Indeed, it was not obvious in advance that this would be the result, given that the temporal fine structure of TFS_Story was so heavily modulated by the envelope of Env_Story for the chimeras. Moreover, this pattern cannot be because the Re-Env was correlated with the raw TFS_Story envelope, because, as mentioned above, if anything, they are weakly negatively correlated with each other (Table 1 – second column). As such, it appears that this result supports that notion that cortex has been able to synthesize a coherent representation of the TFS_Story from its (partial) temporal fine structure. However, just because the pattern of TFS_Story results appears qualitatively similar for the raw envelope analysis (Fig 3*B*) and the Re-Env analysis (Fig 3*C*), does not allow us to conclude that the signal being tapped into by the two analyses is actually the same. To explore this, we directly compared the stimulus reconstructions that were obtained using the raw envelope of the TFS_Story with those obtained using the envelope recovered from the TFS structure of the chimera. There was a significant positive correlation between these reconstructions only for the 1-band condition (*r*_*s*_ = 0.18, *p* < 0.001, 95% CI [0.16-0.20]; Table 1 – right column). And, while this correlation was relatively low, it was substantially higher than the correlations between the reconstructions and the original or reconstructed envelopes shown in Figs 3B and 3C. This suggests that the information being recovered by cortex – as indexed by the Re-Env based reconstruction analysis – is not identical to that being indexed by the raw envelope analysis, but that there is some overlap between the two. Again, it is not surprising that they are not identical given that the temporal fine structure of TFS_Story was so heavily modulated by the envelope of Env_Story for the chimeras. And, again, this is the reason why Env_Story reconstructions based on the raw and chimera envelopes are so highly correlated with each other (Table 1 – third column). A final observation is the correspondence between the behavior and speech reconstructions for TFS_Story in the 16-band condition. At the group level, behavioral performance was not above chance, indicating a general lack of understanding of TFS_Story in this condition, consistent with previous research (Smith et al., 2002). This was mirrored in the failure to reconstruct the recovered envelope of TFS_Story in this condition (Fig. 3*C*).

**Table 1.** Stimulus and Reconstructed Envelope Correlations. Computed between Raw and Chimera derived envelopes for both the original Stimulus and TRF reconstructed versions.

|  | Stimulus |  | TRF Reconstruction |  |
| --- | --- | --- | --- | --- |
|  | Env_Story | TFS_Story | Env_Story | TFS_Story |
|  | Raw - Chimera | Raw - Chimera | Raw - Chimera | Raw - Chimera |
| 1-nBands | 0.91 (0.001) | -0.20 (0.001) | 0.91 (0.001) | 0.18 (0.002) |
| 4-nBands | 0.90 (0.001) | -0.11 (0.001) | 0.97 (0.001) | 0.08 (0.114) |
| 16-nbands | 0.73 (0.001) | -0.10 (0.001) | 0.95 (0.001) | -0.10 (0.019) |

Notes: Statistics reported as: Pearson's rho (*p*-value). Correlations computed using Robust Correlation Analyses with Bootstrap resampling across subjects (Pernet et al., 2013).

### Intelligibility of speech-speech chimeras is reflected more strongly by cortical speech tracking in the delta-band than the theta-band

A substantial amount of previous research has sought to distinguish the roles of delta-and theta-band cortical activity in speech processing (Nai Ding & Simon, 2013, 2014; Doelling et al., 2014; Etard & Reichenbach, 2019; J. E. Peelle, J. Gross, & M. H. Davis, 2013). With that in mind, we sought to explore the sensitivity of delta and theta band speech tracking measures across our different chimera conditions. We did this by attempting to reconstruct the envelopes of Env_Story and TFS_Story from EEG that had first been filtered into either the delta (0.05–4 Hz) or theta (4-8 Hz) bands (using zero phase-shift Chebyshev type-2 bandpass filters). We found that the pattern of reconstruction accuracies based on delta band EEG (Fig. 4, left panel) corresponded to that seen for speech intelligibility (Fig. 3*A*). However, this was much less clear for reconstructions based on theta band activity (Fig. 4, right panel).

**Figure 4.**
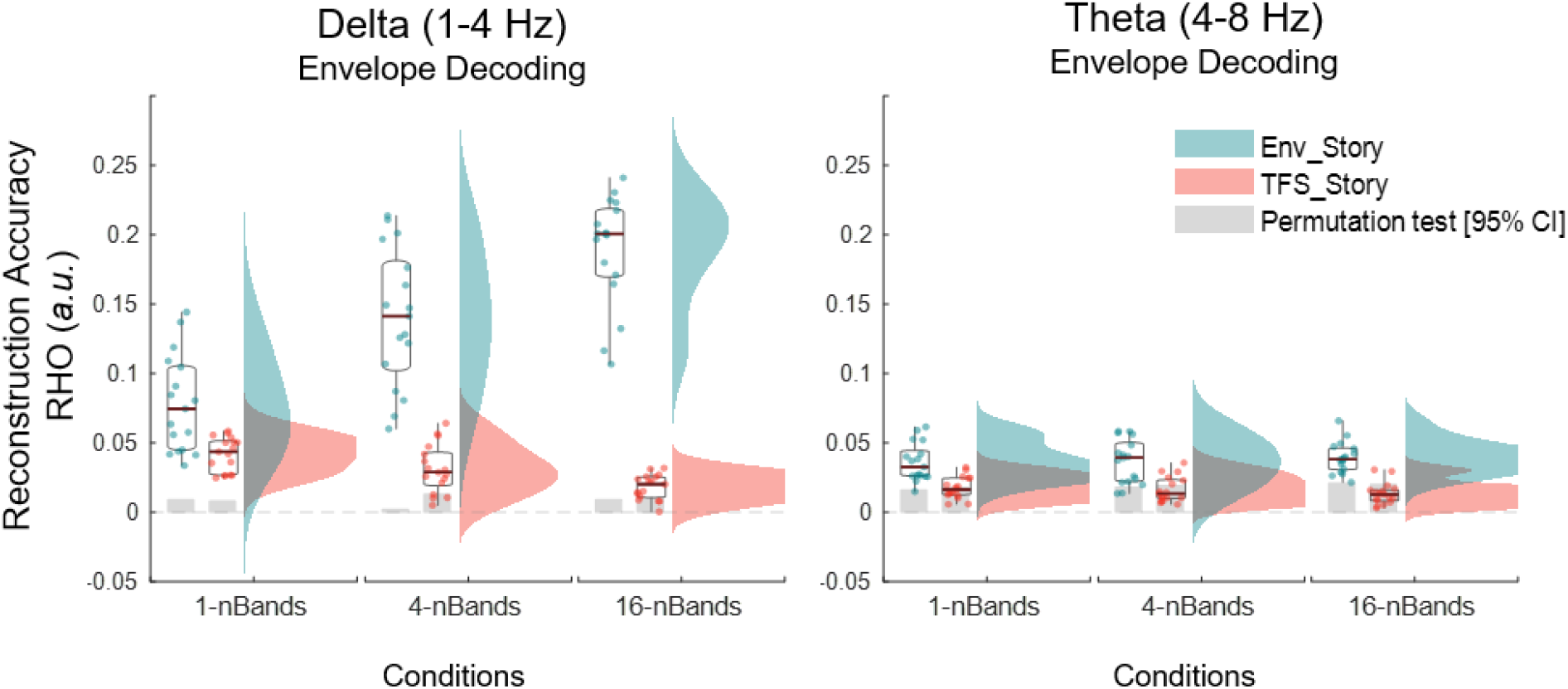
Envelope reconstructions based on delta and theta band EEG. Box plots (mean ± SEM) and rain kernel density estimates of reconstruction decoding accuracy (Rho) values for Env_Story (turquoise) and TFS_Story (peach). Group- and individual statistics; black line in box plot indicates mean across subjects, asterisks indicate group statistical significance (permutation test, *p < 0.05) and grey box demarcates single subject-level statistical significance above chance (permutation test, *p < 0.05). Single dots represent single subject data.

Indeed, for delta band reconstructions, as with both the broadband EEG decoding of the envelopes of the original speech (Fig. 3*B*) and the chimera speech (Fig. 3*C*), we found significant main effects for *chimera condition* (*F*_(1,2)_ = 45.12, *p* < 0.001, η^2^ = 0.58, *BF*_10_ = 6.01) and *story* (*F*_(1,1)_ = 34.44, *p* < 0.001, η^2^ = 0.16, *BF*_10_ = 4.287). And, we again saw a significant two-way interaction between *chimera condition* and *story* (*F*_(1,2)_ = 132.42, *p* < 0.001, η^2^ = 0.15, *BF*_10_ = 20.08). Again, post-hoc Bonferroni-corrected pairwise *t*-tests revealed that reconstructions for Env_Story were significantly higher for the 16-band condition than 1-band condition (*t*_(16)_ = 16.9, *p* < 0.001, *BF*_10_ = 8.88 ×10 ^8^), and TFS_Story reconstructions were significantly higher for the 1-band condition than the 16-band condition (*t*_(16)_ = 5.14, *p* < 0.001, *BF*_10_ = 283.93), as we had found with broadband EEG. In contrast, theta-based reconstructions did not track significantly with speech intelligibility in the chimera stimuli. While there was a main effect for *story* (*F*_(1,1)_ = 92.12, *p* < 0.001, η^2^ = 0.5, *BF*_10_ = 4.84), there was no main effect for *chimera condition* (*F*_(1,2)_ = 0.04, *p* = 0.9, η^2^ = 1.11 ×10^−15^, *BF*_10_ = 0.002), and no interaction effect (*F*_(1,2)_ = 1.89, *p* = 1.89, η^2^ = 0.01, *BF*_10_ = 0.06). Post-hoc Bonferroni-corrected *t*-tests showed that Env_Story reconstruction accuracies for the 16-band condition were not significantly different (or higher) than the 1-band condition (*t*_(16)_ = 1.2, *p* = 0.24 (*ns*), *BF*_10_ = 0). Similarly, there was no significant difference between 1-band and 16-bands conditions in TFS_Story reconstruction accuracies *t*_(16)_ = 1.94, *p* = 0.07, *BF*_10_ = 1.1).

### Exploring the relationship between the intelligibility of speech - speech chimeras and cortical speech tracking at different latencies

Previous research has suggested that different hierarchical stages of acoustic and linguistic speech processing can be approximately indexed by exploring brain responses to speech at different latencies (Salmelin, 2007). Given that we are interested in disambiguating general acoustic processing from speech-specific processing, we sought to explore the possibility that speech tracking at different time lags might be differentially sensitive to the intelligibility of our various speech chimeras. To do this, we plotted the (forward) temporal response function (TRF), by learning a linear mapping from the envelope of our different chimera stimuli to the corresponding EEG responses (Crosse et al., 2016). If acoustic processing is indexed at shorter latencies and speech-specific processing at longer latencies, we might expect to see strong Env_Story TRF responses at short latencies for all conditions, with variations in longer latency TRF components that mirror our behavioral measures of speech intelligibility. And we might expect to see relatively weak early TRF responses to the TFS_Story with longer latency TRF components again reflecting our speech intelligibility measures.

Figure 5*A* displays TRFs averaged over 12 frontotemporal channels (6 from the left scalp and their symmetrical counterparts on the right), chosen based on previous literature (M. P. Broderick, A. J. Anderson, G. M. Di Liberto, M. J. Crosse, & E. C. Lalor, 2018; M. J. Crosse et al., 2016; Di Liberto, O’Sullivan, & Lalor, 2015; Teoh & Lalor, 2020). Robust TRFs were visible for most conditions. In general, the TRFs for the Env_Story appeared to increase in amplitude from the 1-band condition to the 16-band condition. However, there was no clear difference in the pattern across conditions for early TRF components compared to later components. Indeed, Monte Carlo cluster-based permutation statistics show main effects of chimera condition at a number of latencies both early and late (subplot Fig. 5). For the TFS_Story, the 1-band (most intelligible) condition displayed the largest TRF at a latency of ∼100-170 ms. However, this result needs to be treated with caution given that the (somewhat intelligible) 4-band response appeared smaller than the (completely unintelligible) 16-band response.

**Figure 5.**
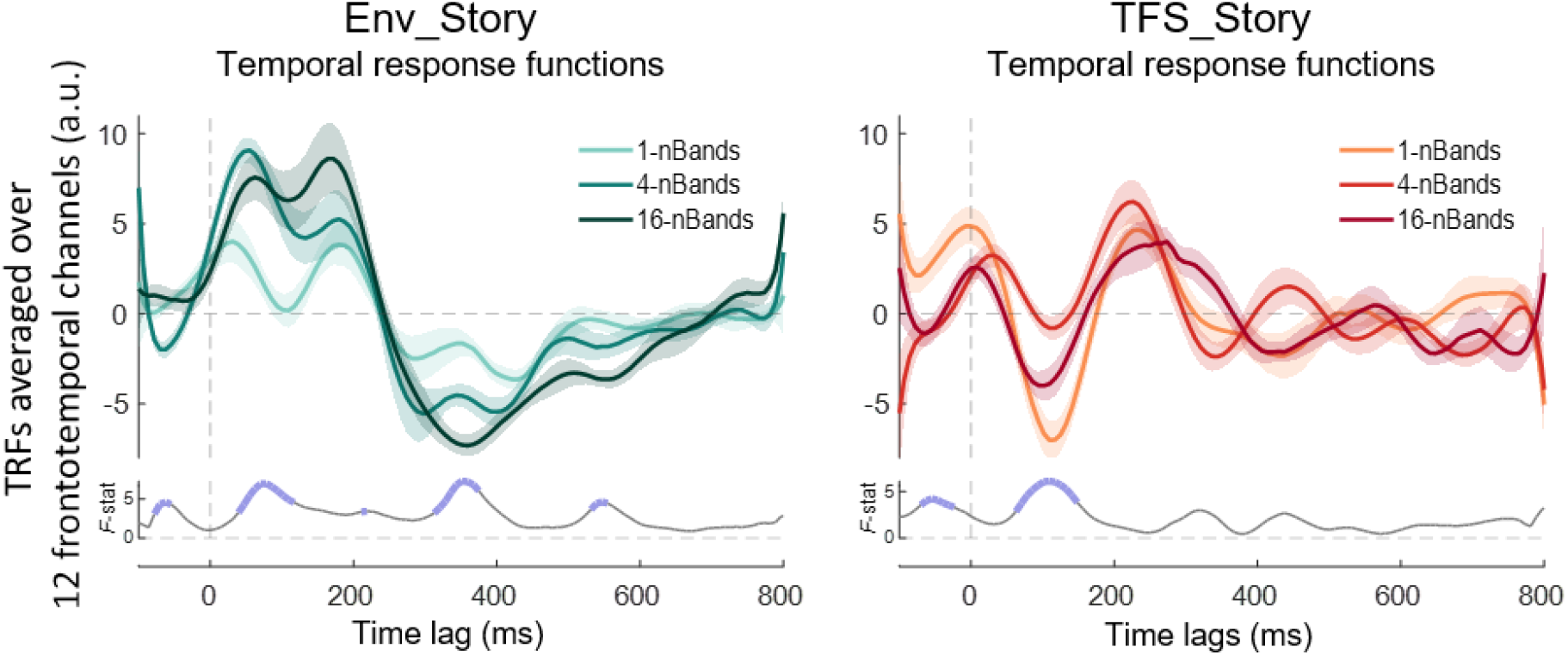
Temporal response functions (TRFs) indexing the relationship between the envelope of the different chimera stimuli and the corresponding EEG response. TRFs are averaged over a set of 12 electrodes with high prediction accuracies over frontotemporal scalp (6 on the left side of the scalp, and their symmetrical counterparts on the right), without biasing any of the TRF models. Shaded lines demarcate SEM. Subplots show Monte Carlo cluster-based permutation statistics, main effect of chimera condition (F-value), with the thick blue lines indicating significance (*p* < 0.05).

### Modeling speech responses in terms of both acoustics and phonetic features can disambiguate acoustic and speech - specific processing

The envelope reconstruction results above (Fig. 3) highlight one of the key limitations of using envelope tracking as a measure of speech processing. The general correspondence in the patterns of results for behavior and tracking indicate a sensitivity to speech-related processing. However, the robust tracking of speech that is either completely or almost completely unintelligible (i.e., Env_Story in the 1-band and 4-band conditions; Fig. 3*B*), highlights that much of the cortical tracking is simply driven by acoustic energy – consistent with previous work (Howard & Poeppel, 2010; Lalor et al., 2009). In the previous section, we attempted to approximately index EEG activity that is driven by the stimulus acoustics and that which reflects speech-specific processing by exploring TRF components at different latencies. However, that analysis is still based on a very coarse representation of the stimulus (i.e., its envelope) whose dynamics necessarily correlate with those of many acoustic and linguistic features. And, as such, the results are difficult to interpret.

An alternative approach is to use forward encoding models to explicitly model EEG responses in terms of specific acoustic and linguistic features of the speech stimulus (Di Liberto et al., 2015). Here we explore this issue with a focus on acoustic and phonemic features whose dynamics likely correlate with those of the envelope. In particular, we aimed to model EEG responses in terms of both the spectrogram and phonemes of speech, and to try to identify unique variance in the EEG responses that can be explained by each. In doing so, we aim to disentangle the encoding of acoustic features from speech-specific processing that likely jointly contribute to the envelope reconstruction measures investigated above. Specifically, there are redundancies between the predications of feature spaces, so here we were interested in clearly isolating unique contributions of each feature.

To explicitly quantify the unique contributions of acoustic and phonemic processing to the EEG, we computed the partial correlation coefficients (Pearson’s *r*) between the EEG predicted by either the Phoneme (*Ph*) or Spectrogram (*Sgram*) model with the actual recorded EEG, after controlling for the effects of the other feature. The unique predictive power of each model is shown in Fig. 6 for phonemes and spectrogram across each band condition (shown here for an average across 12 channels, six symmetric pairs on the left and right scalp as previously used in Di Liberto et al., 2015, 2018, although including all 128 channels revealed the same qualitative pattern of results). In particular, we conducted two separate ANOVAs, one focused on *Ph* predictions (whilst controlling for *Sgram*) and one focused on *Sgram* (whilst controlling for *Ph*). Both ANOVAs (*rm*ANOVA) had factors of *chimera condition*; 1-, 4-, 16-bands, and *story:* Env_Story, TFS_Story). Both models showed a generally similar pattern to our envelope-based results above, with improvements in prediction with increasing chimera bands for Env_Story and decreases in prediction with increasing chimera bands for TFS_Story (Fig 3). Indeed, for both models, we found a significant main effect of *chimera* c*ondition* (*Ph*: (*F*_(1,2)_ = 3.06, *p* < 0.05, η^2^ = 0.18, *BF*_10_ = 59.3); *Sgram*: (*F*_(1,2)_ = 5.55, *p* = 0.006, η^2^ = 0.09, *BF*_10_ = 3.2 ×10 ^17^)), and *Story* (*Ph*: (*F*_(1,1)_ = 32.53, *p* < 0.001, η^2^ = 0.41, *BF*_10_ = 1.61 ×10 ^08^); *Sgram*: (*F*_(1,1)_ = 62.89, *p* = 0.001, η^2^ = 0.53, *BF*_10_ = 1.38 ×10 ^17^)), and a significant interaction of *chimera condition* and *story* (*Ph*: (*F*_(1,2)_ = 16.1, *p* = 0.001, η^2^ = 0.34, *BF*_10_ = 267.65); *Sgram*: (*F*_(1,2)_ = 21.98, *p* < 0.001, η^2^ = 0.07, *BF*_10_ = 2.9 ×10^16^)). Additionally, post-hoc Bonferroni-corrected pairwise *t*-tests revealed that Env_Story partial correlation coefficient scores in the 16-band condition were significantly higher than those scores in the 1-band condition for both models (*Ph*: (*t*_(16)_ = 4.65, *p* < 0.001, *BF*_10_ = 120.12); *Sgram*: (*t*_(16)_ = 5.44, *p* < 0.001, *BF*_10_ = 481.9)), and TFS_Story scores in the 1-band were significantly higher than the 16-bands for both models (*Ph*: (*t*_(16)_ = 2.79, *p* < 0.001, *BF*_10_ = 4.28); *Sgram*: (*t*_(16)_ = 3.78, *p* < 0.05, *BF*_10_ = 23.68).

**Figure 6.**
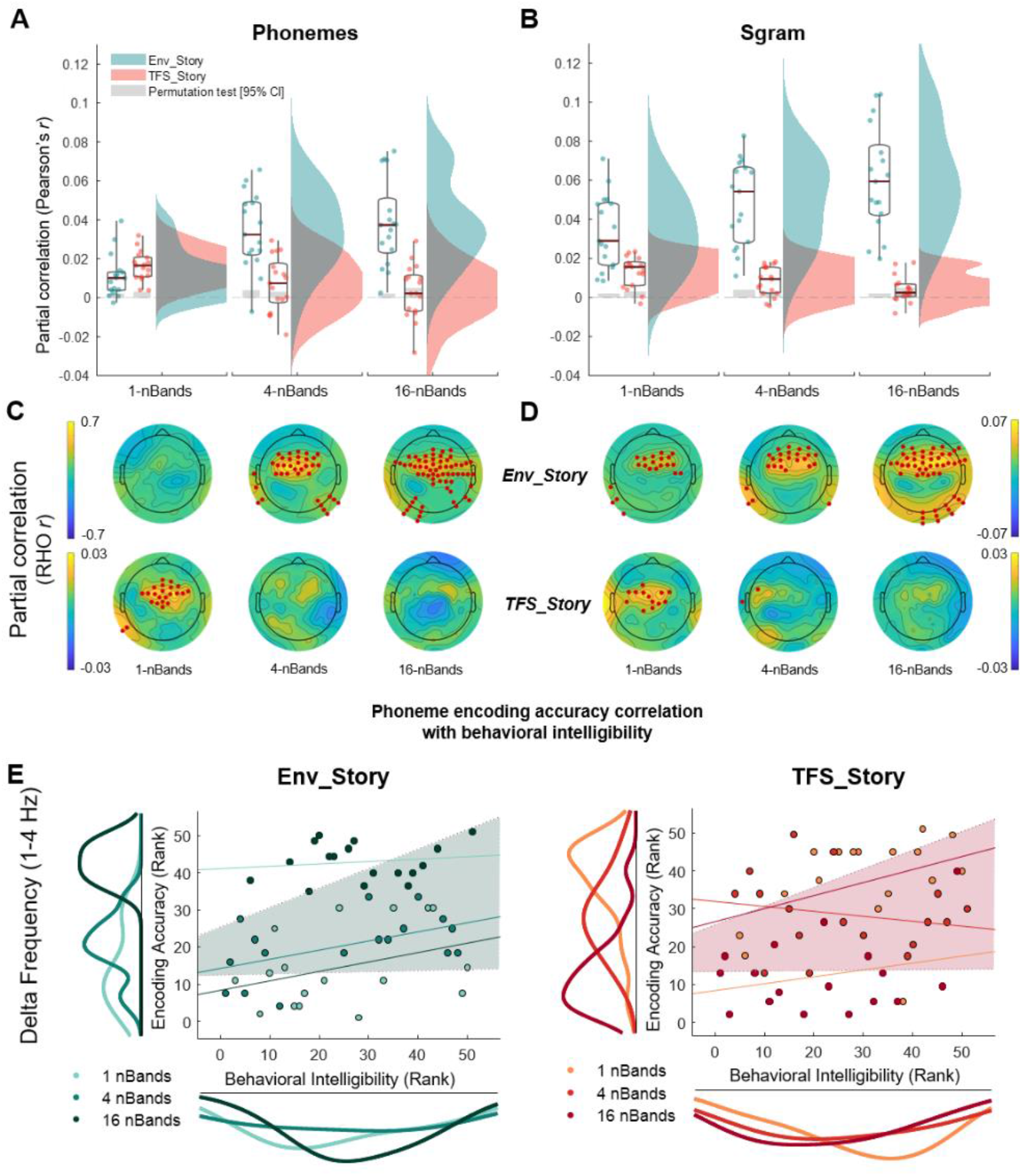
Partial correlations of real EEG and EEG predicted using forward encoding models based on different stimulus feature representations. **A**. Partial correlation of real EEG and EEG predicted using the phonetic feature representation (Phon) while controlling for the acoustic representation (Sgram). Correlations are averaged over the same set of 12 electrodes as Fig. 5, i.e., a set with high prediction accuracies over frontotemporal scalp (6 on the left side of the scalp, and their symmetrical counterparts on the right). **B**. Partial correlation of real EEG and EEG predicted using the Sgram representation while controlling for the Phon representation. Box plots (mean ± SEM) and rain kernel density estimates of Partial correlation Rho (Pearson’s r, two-tailed) values for Env_Story (turquoise) and TFS_Story (peach). Group-and individual statistics; black line in box plot indicates mean across subjects, asterisks indicate group statistical significance (permutation test, *p < 0.05) and grey box demarcates single subject-level statistical significance above chance (permutation test, *p < 0.05). Single dots represent single subject data. **C, D** Topographical plots show partial correlation Rho across channels for Env_Story (**C**) and TFS_Story (**D**). Red dot indicates significant effects at group level statistics (one-tailed cluster-based permutation test, *p< 0.05). **E**. Phoneme encoding according to accuracy relationship with behavioral intelligibility scores for delta (1-4 Hz) frequency band. **Left**, Env_Story and **Right**, TFS_Story. Individual dots represent subjects and are colour coded according to condition. Using Robust Spearman’s correlations (bootstrap permutation test p < 0.05, confidence interval 95%). This was done for each individual condition (colored lines), as well as collapsed across all conditions (shaded area representing confidence interval 95%). Subplots show the distribution of the data in terms of both envelope decoding (left) and behavior (below) using the same color code as the subject dots.

While these data show that both acoustic and phonemic features track with our original envelope findings, there are a couple of notable differences in the performances of the two models. First, in the 1-band condition, the Env_Story is completely unintelligible, so any prediction based on Env_Story must be driven by acoustic processing. This is reflected in the fact that the acoustic *Sgram* model (Fig 6*B,D*) predictions are significantly above zero (*z* = 3.38, *p* = 7.13 ×10 ^−04^). Meanwhile, the unique contribution of the *Ph* model (Fig 6*A,C*), is not significantly above zero for this condition (*z* = −0.4, *p* = 0.71), which again makes sense given that there is no phonemic information in this chimera for Env_Story as speech was degraded and unintelligible in this band. Conversely, for TFS_Story in the 1-band condition, the *Ph* model adds unique predictive power (Fig 6*A,C*), again, reflecting the fact that this chimera contains phonemic information for TFS_Story. The unique contribution of the *Ph* model disappears for the 4-band and 16-band conditions, again supporting the idea that it represents a measure of speech-specific processing, separable from more general acoustic processing. The differences between the *Ph* and *Sgram* models can also be seen when visualizing which EEG channels are significantly predicted by each model (Fig 6 *C,D*). While the performance of the *Sgram* model is qualitatively similar to the envelope, the unique contributions of the *Ph* model seem to track more specifically with speech intelligibility.

Next, under the assumption that the phonetic feature model performance might more closely relate to speech intelligibility, we assessed the relationship between EEG phonetic feature encoding (while controlling for Sgram) and behavioral intelligibility score (Fig. 6*E*). In particular, we ran a Spearman’s Robust correlation analysis with bootstrap resampling (Pernet et al., 2013), and specifically looked at delta band EEG (given our results in Fig. 4). As before, we carried out the analysis in one correlation collapsing across conditions and found a significant positive correlation between phonetic encoding and behavioural intelligibility score as a function of condition, for Env_Story (*r*_*s*_ = 0.46, *p* < 0.001, 95% *CI* [0.17 0.68]) and TFS_Story (*r*_*s*_ = 0.36, *p* < 0.05, 95% *CI* [0.11 0.58]). As we collapsed across conditions, we wanted to provide further support for this relationship using a linear-mixed effect model (LME) with behavioral intelligibility score and phonetic feature encoding as dependent and predictor variables respectively. The LME models variability due to stimulus conditions and subjects simultaneously. We found a significant positive relationship between reconstruction accuracy and intelligibility score for Env_Story (*β* = 302.65, *SE* = 101.56 *p* < 0.001) and for TFS_Story (*β* = 154.33, *SE* = 62.12 *p* < 0.01). Non-significant within-condition correlation lines are plotted for completeness.

## 6. Discussion

In this study, we have aimed to disentangle the contributions to cortical speech envelope tracking that derive from acoustic and speech-specific processing. Using speech-speech chimeras, we aimed to decouple the acoustic envelope fluctuations of a stimulus from its speech content. We found evidence of robust envelope tracking based on acoustic energy fluctuations. We also found that changes in the strength of envelope tracking correlate with speech intelligibility across conditions. Altogether, we conclude that the cortical tracking of speech envelopes contains a large contribution from general auditory processing, with additional contributions from neural populations that are specifically tuned to speech, consistent with our original hypothesis (hypothesis 4, Fig. 2).

The notion of a substantial general acoustic contribution to cortical speech responses is in line with the idea that speech sounds are perceived using mechanisms that evolved to process environmental sounds more generally (Diehl et al., 2004), with additional linguistic processing occurring in specialized downstream pathways (Hickok & Poeppel, 2007; Rauschecker & Scott, 2009). Indeed, there is a wealth of evidence suggesting that speech is processed by a hierarchically organized network of cortical regions with responses in earlier stages (including primary auditory cortex) being well accounted for based on the spectrotemporal acoustics of the stimulus, and later stages being invariant to those acoustics and involved in more abstract linguistic processing (Davis & Johnsrude, 2003; de Heer et al., 2017; DeWitt & Rauschecker, 2012; Huth et al., 2016; Kell et al., 2018; Norman-Haignere & McDermott, 2018). Of course, to contribute to envelope tracking, any such linguistic processing must involve speech features whose dynamics correlate with the envelope. One candidate set of features would be phonemes, whose onsets and offsets will often coincide with fluctuations in the envelope. The superior temporal gyrus – an auditory association area whose activity is not especially well captured based on a spectrotemporal representation of speech (Davis & Johnsrude, 2003; Norman-Haignere & McDermott, 2018) has shown a high degree of tuning for phonetic features (Chang et al., 2010; Mesgarani, Cheung, Johnson, & Chang, 2014).

The idea that envelope tracking consists of general acoustic processing contributions from primary auditory areas, and temporally correlated speech-specific contributions from areas like STG explains many of the features of our results. For example, we see robust envelope tracking of Env_Story in the 1-band condition, when it is completely unintelligible. This must necessarily be due to simple auditory responses to changes in acoustic energy and fits with previous work showing robust envelope tracking to non-speech stimuli (e.g., Lalor et al., 2009). The tracking of the Env_Story envelope then increases in strength across the 4-band and 16-band conditions as subject increasingly understand the speech content of story 1, which likely reflects the increased contribution from speech-specific neuronal populations across these conditions. Conversely, we see significantly lower envelope tracking for TFS_Story, which makes sense as the energy fluctuations of the stimulus are dominated by Env_Story. Thus, the general acoustic processing contribution will be relatively insensitive to the dynamics of TFS_Story. However, in the 1-band (and 4-band) condition, subjects can partially understand the speech content of TFS_Story. And this leads to significant envelope tracking for TFS_Story in that condition, likely driven by contributions from STG and other speech-specific areas. That said, we also found that the EEG tracked the envelope that was recovered from the chimera stimuli (RE-Env), meaning that there could be tracking of lower-level acoustic features that have been resynthesized from the chimera after it passes through the cochlea.

We attempted to parse the contributions of acoustic and speech-specific processing using forward encoding models. We did this in two ways. First, we explored whether or not we might see differential sensitivity to variations in intelligibility as a function of latency between stimulus and response (Fig. 4). While no particularly clear pattern emerged, it is important to bear in mind that the envelope of speech is a very compressed measure of a speech signal. Indeed, any speech features (acoustic or linguistic) that covary with the speech envelope are likely to contribute to the output measure. As such, the TRF in this case is very difficult to interpret. Second, we explored how the EEG across our different conditions reflected acoustic and phonetic feature processing by explicitly modeling those features. In general, both sets of features contributed uniquely to predicting EEG responses, with both increasing across conditions for Env_Story and decreasing across conditions for TFS_Story. Notably, the unique contribution for the phoneme features (*Ph*) was only significant for individual EEG channels for Env_Story in the 4-band and 16-band conditions, i.e., where Env_Story could be partially understood, and TFS_Story in the 1-band condition, i.e., were TFS_Story could be partially understood. The phoneme model added no value in conditions where Env_Story (1-band) and TFS_Story (16-band) could not be understood. This suggests that modeling the responses to speech in this way has enabled us to tap into contributions from speech-specific areas, like STG.

The idea of general and speech-specific contributions to envelope tracking helps to explain why it can be quite difficult to link cortical envelope tracking measures with measures of speech understanding (Howard & Poeppel, 2010). For example, some studies have reported that cortical envelope tracking shows sensitivity to speech intelligibility (Jonathan E Peelle et al., 2013; Vanthornhout, Decruy, Wouters, Simon, & Francart, 2018), while others have failed to find it (Howard & Poeppel, 2010). It may be that the large general auditory processing contribution to envelope tracking that is common to both intelligible and unintelligible speech somewhat masks a smaller contribution from task-based speech-specific processing. Indeed, in studies that have tried to control or account for the acoustic contributions to speech perception, correlations with behavior have been reported for phoneme-level processing (Di Liberto et al., 2018), including in STG (Leonard et al., 2016).

It has been well established that cortical envelope tracking is strongly affected by selective attention (Nai Ding & Simon, 2012; Kerlin, Shahin, & Miller, 2010; O’Sullivan et al., 2015; Power et al., 2012). However, correlations between envelope tracking and behavioral measures of cocktail party attention have been difficult to identify (O’Sullivan et al., 2015; Tune, Alavash, Fiedler, & Obleser, 2020). Again, if we consider envelope tracking as the combination of general acoustic and speech-specific processing, this makes sense. Using invasive recordings, it has recently been shown that cocktail party attention effects vary substantially in their strength across the cortical hierarchy, with weak effects in early auditory areas like Heschl’s gyrus and much stronger effects in areas like STG (O’Sullivan et al., 2019). And EEG studies have suggested something similar on the basis of examining envelope tracking at different latencies (Power et al., 2012) or based on trying to isolate markers of acoustic and phonetic feature processing (Teoh & Lalor, 2020). Indeed, focusing specifically on cortical responses to higher-level linguistic speech features – including those at the lexical and semantic levels – researchers have often found strong correlations with attention (Brodbeck, Hong, & Simon, 2018; Michael P Broderick, Andrew J Anderson, Giovanni M Di Liberto, Michael J Crosse, & Edmund C Lalor, 2018), and speech understanding more generally (Michael P Broderick et al., 2018). Incidentally, it is worth reflecting on the possible role of attention in the pattern of results we see in the present study. For example, one might wonder if changes in the amount of attention being paid to Env_Story or TFS_Story across conditions is driving the changes in envelope reconstruction accuracy we see (Fig 2). We think this is unlikely. This is because, in our experiment, there was always just a single chimera stimulus being presented at any one time. Thus, any increase in attention to the speech content is likely to enhance the response to the acoustic energy changes. Of course, this is not guaranteed. Feature-based attention has been shown to affect the processing of some features more than others within the same object (Maunsell & Treue, 2006). However, the limited literature exploring how different feature-based attention tasks within a cocktail party environment have shown no clear effects on envelope tracking (Lauteslager, O’Sullivan, Reilly, & Lalor, 2014).

The approach and results in this study have implications for theories of so-called speech entrainment (Giraud & Poeppel, 2012; Obleser & Kayser, 2019). One such prominent theory posits that intrinsic, ongoing oscillatory brain rhythms “entrain” to the rhythms of the speech signal by aligning their phase with the stimulus in an anticipatory, behaviorally effective manner. The core idea of this model is that salient points (‘edges’) in the speech signal cause “phase resetting” of ongoing oscillatory activity. The realignment of the phase of these oscillations, which has been linked to fluctuations in cortical excitability (Lakatos et al., 2005), then enables the parsing and chunking of the continuous speech signal into discrete linguistic units for further processing by faster oscillations (Ghitza, 2011; Giraud & Poeppel, 2012; Rimmele, Morillon, Poeppel, & Arnal, 2018). This theory links to fundamental electrophysiological observations in nonhuman studies (Lakatos, Karmos, Mehta, Ulbert, & Schroeder, 2008). However, when it comes to the specific case of speech processing in humans, it has largely been built from observations that cortical activity (often measured noninvasively) shows consistent phasic fluctuations in theta band activity across repeated presentations of a speech stimulus (Huan Luo & Poeppel, 2007). The fact that theta band activity appears special in this regard has led to the suggestion that this entrainment facilitates the parsing of continuous speech into syllables, given that the timescale of syllables is in the range of 4-8 Hz. However, basing this theory on the idea of consistent fluctuations in noninvasively recorded brain responses to speech is problematic. As we have shown in the present study, much of the variance in these fluctuations is driven by amplitude modulations of the stimulus, again, in line with previous research (Lalor et al., 2009). If acoustic edges (perhaps corresponding to syllable boundaries) were driving this cortical tracking, we might expect to see stronger tracking for TFS_Story than Env_Story in our 1-band condition. Of course, it may be that different low-frequency oscillators, each with a different specific role, are concurrently active in different early cortical areas. Indeed, recent research (also using stimuli where the ENV and TFS have been decoupled) has reported cortical entrainment in the theta range to temporal fine structure, distinct from that to the envelope (Teng, Cogan, & Poeppel, 2019). In particular, that study showed robust MEG tracking of unintelligible stimuli composed entirely of the TFS of speech (i.e., with no envelope fluctuations). Furthermore, behavioral results showed that TFS could help improve the recognition of speech whose envelope had been temporally distorted. The authors interpreted these findings as evidence that cortical entrainment to speech reflects the tracking of both the temporal and spectral structure of speech. That finding agrees well with our data showing EEG tracking of both Env_Story and TFS_Story. However, differences in the stimuli and tasks between the two studies led to different interpretations of the results. Teng et al. (2019) interpret their findings as evidence for two complementary mechanisms through which neural entrainment can facilitate the segmentation of speech (for further processing). On the other hand, our somewhat more direct linking of envelope and TFS tracking to speech comprehension in one experiment has led us to the interpretation that these neural measures index, not just the segmentation of speech, but dissociable contributions from general acoustic and speech-specific processing. Specifically, our interpretation is that acoustic energy fluctuations drive evoked responses in early auditory areas, with speech-specific tuning leading to additional contributions from auditory association areas like STG. We think this interpretation can also explain the cortical tracking of TFS reported by Teng et al. (2019). Although decisively adjudicating on the relative contributions of entrained oscillations vs evoked responses is not straightforward (Obleser & Kayser, 2019).

Finally, the variation in speech intelligibility across chimera conditions was reflected more strongly in delta band EEG frequencies than theta band. This agrees with previous studies highlighting a specific correspondence between speech intelligibility/comprehension in challenging listening environments and delta band tracking (Ding et al., 2014; Etard & Reichenbach, 2019(Mai & Wang, 2019)). For example, Etard & Reichenbach (2019) explored EEG responses to native and foreign language in different levels of background noise and found that cortical tracking in the theta band was mainly correlated to clarity, whereas the delta band was most closely related to speech comprehension. And Ding et al., (2014), who dissociated the envelope and temporal fine structure of speech using noise vocoding, found that cortical tracking in the delta band predicted speech recognition scores for individual listeners. Indeed, more generally, a slew of recent research has linked cortical activity in the delta band with the tracking of linguistic features, even if these features have no acoustic correlates (Brodbeck et al., 2018; Broderick et al., 2018; Buiatti et al., 2009; Ding et al., 2016a; Jin et al., 2018; Makov et al., 2017; Sheng et al., 2019).

While the correspondence of delta band activity to behavior was not unexpected, we were somewhat surprised that the delta band tracking of the chimera stimuli was generally so much stronger than for the theta band. We sought to understand this through the lens of our previously mentioned interpretation that cortical speech tracking is largely driven by evoked responses. To that end, we calculated the mean amplitude of the modulation spectrum of our chimera stimuli in the delta and theta frequency ranges, and found that delta-band stimulus modulations were in excess of three times as large as modulations in the theta range (1-band: 3.15; 4-band 3.39; 16-band: 3.34). Thus, the stronger tracking of speech in the delta band may simply reflect differences in evoked response amplitude based on more energetic modulations in that frequency range. Even if entrained oscillations are playing an important role in our data, the larger delta band responses would still fit with some previous studies. For example, when the spectral resolution of speech decreases, theta-band cortical tracking has been shown to be reduced (Peelle et al., 2013; Ding et al., 2014), but delta-band tracking can be enhanced (Ding et al., 2014). Furthermore, when speech is presented in competing speech streams, delta-band frequency (1-4 Hz) cortical tracking has been reported to be robust, while theta-frequency band (4-8 Hz) cortical tracking often decreases as the level of the competing stream is increased (Nai Ding & Simon, 2013). That said, we think the characteristics of the stimulus modulation spectrum remains the best explanation, especially of the relatively poor theta-band envelope tracking of some of the more intelligible chimera stimuli (i.e., the 16-band stimulus). Further work is needed to more fully explore the roles of evoked responses and entrained oscillations in the delta and theta bands, bearing in mind that both general auditory and speech-specific activity are likely to be occurring within those frequency ranges. In any case, it is clear that continued future efforts at understanding speech processing must account for brain responses that reflect general acoustic processing.

